# The phosphoS655 Alzheimer’s Amyloid Precursor Protein (APP) interactome in neuronal differentiation

**DOI:** 10.64898/2026.03.26.714600

**Authors:** David L. Almeida, Joana F. da Rocha, BC da Cruz, J. Mirjam A. Damen, A. F. Maarten Altelaar, H. Osório, Odete A. B. da Cruz e Silva, Sandra I. Vieira

## Abstract

The Alzheimer’s Amyloid Precursor Protein (APP) has determinant roles in neuronal development and function, both in its full-length conformation and as some of its proteolytic peptides, particularly secreted (s)APPa. Given that APP phosphorylation tightly regulates its trafficking, proteolysis, and protein-protein binding, it consequently affects several APP functions. The S655 residue, located in the basolateral sorting motif YTSI at APP C-terminus has been observed to be phosphorylated in mature full-length APP and its C-terminal fragments. Previously observed to modify APP’s protein interactions, resulting in altered endolysosomal trafficking, andincreased half-life and sAPPa generation, phosphoS655 APP has potential to modulate APP-mediated neuronal differentiation. To study the phosphoS655 differential interactome relevant for neuronal differentiation, SH-SY5Y cells expressing Wt or S655 phosphomutants APP-GFP were differentiated at two time points. APP-GFP and their respective interacting partners were immunoprecipitated using GFP-trap, and interactors identified by mass spectrometry. Both dephospho and phosphoS655 interactomes were generally enriched in similar processes, primarily RNA processing and translation, as well as signal transduction, metabolism, and cytoskeleton remodeling. The smaller phosphoS655 interactome contributes for functional specialization via binding to e.g. FUBP3, ELAVL4, ATXN2, Tubulin, INA. Several of these specific binding partners are known to promote neurite outgrowth and likely underlie our experimental observation that phosphoS655 APP promotes neuritogenesis, particularly the formation of longer neuritic extensions. These results are not only important for the body of knowledge on this Alzheimer’s disease core protein, but may also aid in future therapies against this disease.

## Introduction

The Amyloid Precursor Protein (APP) is mainly studied for its association with Alzheimer’s disease (AD). This transmembrane protein is dynamically sorted through the membranes of the cell and intracellular organelles. During trafficking, full-length APP is cleaved by proteases into some functional peptides. Briefly, α-secretases generate sAPPα and a membrane-tethered C-terminal fragment (CTF), whereas cleavage by β-secretase releases the sAPPβ along with a CTF. The resulting CTFs are subsequently processed by the γ-secretase complex to produce the p3 or amyloid-β (Aβ) peptides, respectively, and the intracellular AICD fragment (1,2). In addition to the aforementioned classic APP cleavage pathways, other proteolytic processing events have been described for APP (1).

Several crucial physiological roles have been attributed to the intricate APP molecule, including most recently in maintaining proteostasis via regulation of TGFβ signaling (3); for comprehensive reviews, see (4,5). Particularly, APP has been described to be determinant for neuronal development and function, with attributed functions in cell migration (6), neuritogenesis (7–9) and synaptic structure, transmission, and plasticity (10–12). Strikingly, loss of APP expression due to a homozygous truncating mutation led to developmental delay, microcephaly, callosal dysgenesis, and seizures in a 20-month-old infant (13). Several of these functions have been attributed not only to the full-length protein but also to its proteolytic fragments (14–18). However, contradictory results have been reported regarding their role in nervous system development and function (19–22), and await further investigations.

An additional level of complexity is brought by APP regulation through post-translational modifications. As a phosphoprotein, APP phosphorylation tightly regulates its trafficking (23,24), proteolysis (24,25), and protein-protein binding (24,26). APP CTFs and AICD fragments have also been detected in the phosphorylated form (27). One possible APP phosphorylation site is the S655 residue located in the basolateral sorting motif ^653^YTSI^656^. Phosphorylation at S655 has been detected in mature APP molecules in rat cortex (28), and in APP-CTFs in the hippocampus of AD patients (29). S655 is described to be phosphorylated by protein kinase C and Rho-associated coiled-coil kinase 1 (28,30,31). Nuclear Magnetic Resonance analyses revealed that APP phosphorylation at S655 induces significant local conformational changes within and downstream of the ^653^YTSI^656^ motif (32). These structural alterations likely regulate protein-protein interactions; for instance, the phosphomimetic mutant APP-S655E shows enhanced interaction with the retromer trafficking complex member VPS-35 (23), while APP-CTF-S655E disrupts the interaction with the regulator of lysosomal trafficking AP-3 (33). Ultimately, these alterations in protein-protein interaction directed by S655 phosphorylation lead to enhanced APP half-life and a higher rate of sAPPα secretion, with decreased APP trafficking to lysosomes and reduced Aβ42 production, when compared to the S655A dephosphomimetic mutant (23,33,34).

The present work sought to identify APP binding partners sensitive to its S655 phosphorylation state, a modification that seems to govern protein-protein interactions essential for APP trafficking and processing, and to examine how these interactions change during neuronal differentiation. To this end, we used the neuritogenic cellular model retinoic acid (RA)-differentiated SH-SY5Y cells transiently overexpressing APP-GFP S655 phosphomutants (S655A and S655E). Cells were collected at differentiation days 3 and 7 given that they represent two distinct stages of RA-induced SH-SY5Y neuronal differentiation. In the first one, associated with an increase in the number of pre-neuritic projections, APP levels rise but holo APP is cleaved at a higher rate into sAPP. Neurites appear more abundantly during the second stage, characterized by neuritic elongation and stabilization of longer neurites in a holo APP-assisted manner (35).

The GFPTrap assay was used to immunoprecipitate APP-GFP proteins and their respective interacting partners, which were identified by mass spectrometry (MS). The phophoS655 APP interactome presented a reduced number of protein interactors, when compared to the other categories, denoting a higher degree of functional specificity. In general, the interactors of both S655 dephospho and phosphoAPP were involved in similar biological processes, with some specificities. The phosphoS655 APP interactome presented a special enrichment in some mRNA processing and translation proteins, and in some proteins related to signaling and the cytoskeleton, that are known to interact with each other in a functional network with roles in differentiation, including promotion of neuritogenesis.

## Materials and Methods

### SH-SY5Y cell culture, differentiation and transfection

The SH-SY5Y human neuroblastoma cell line was maintained in minimum essential media/F12 medium supplemented with 10% fetal bovine serum (FBS). Before differentiation, the culture was enriched in the neuroblastic N-type of cells - immature nerve cells that are differentiated with retinoic acid (RA) into neuronal-like cells - based on their lower substrate adherence comparing to the S-type cells (36). N-type enriched cells were plated at 30,000 cells/cm^2^ onto 100 mm culture dishes and incubated at day 0 (D0) with complete medium containing 10 µM RA to start neuronal-like differentiation. Cells were kept in a humidified, 37°C, 5% CO2 incubator and medium was substituted by new medium supplemented with 10 µM RA every other day.

Cells were transfected with TurboFect^TM,^ according to the manufacturer’s instructions (Fermentas Life Sciences), at day 2 or day 6 of differentiation with 9 µg cDNA of human APP isoform 695 wild-type (APPWt), S655A dephosphomutant (APPSA), or S655E constitutive phosphomutant (APPSE), fused with GFP (37). The “empty” GFP vector was used as control. The transfection time points were chosen as key regulatory differentiation periods, based on our previous work on time-dependent APP-induced neuritogenic alterations (36). Transfection complexes were kept for 6h on cells, after which medium was changed. After 24h, the efficiency of cell transfections was confirmed by visualization of GFP expressing cells under an Olympus IX81 epifluorescence microscope, and cells collected.

### GFP Trap IP assays

Pull-down of the GFP moiety of the APP-GFP chimeras using GFP-trap® (Chromotek) according to the manufacturer’s instructions. Upon 24h of transfection, SH-SY5Y cells were washed in PBS, and in 1 mL of ice-cold PBS with PMSF (1:100). Cells were collected by scrapping, centrifuged (5 min, 3000 g, 4°C), and the cells’ pellet lysed for 30 min with 500 µL non-denaturant Lysis buffer (10 mM Tris HCl pH 7.5, 0.5 mM EDTA, 0.5% Gepac-ca-630, 150 mM NaCl, 1 mM PMSF) supplemented with protease inhibitors cocktail (Sigma-Aldrich), 1 mM NaF and 10 mM Sodium Orthovanadate (phosphatase inhibitors). Samples were further centrifuged for 5 min at 20,000 g, an aliquot of supernatant (‘cells lysates’) used to confirm transfection via immunoblot, and the remaining supernatant incubated with previously washed GFP-trap beads (25 µL/sample, 3h with orbital shaking). Beads were magnetically separated until the supernatant was clear and resuspended in wash buffer. Magnetic separation and washing step were repeated 4 times. Beads were resuspended in 100 µL of 2x SDS-sample buffer and boiled for 10 min at 95°C to dissociate immunocomplexes from GFP-trap beads. After vortexed, beads were magnetically separated, and the supernatant transferred to a new microtube. Samples were immediately stored at -20°C for downstream mass spectrometry.

### Mass spectrometry (MS)

For MS analysis, samples were defrosted, 45 μL of each sample were loaded on a 12% SDS-PAGE gel (Bio-Rad 345-0117) and run for 2h under a constant amperage of 20 mA. Afterwards the gel was fixed for 30 min in 40% MeOH, 10% HAc solution and further stained with Coomassie (Gelcode blue stain, Pierce). Each lane was divided in 3 equal gel bands that were excised and cut into small (∼1 mm^3^) pieces. The gel pieces were washed, in-gel reduced with 6.5 mM dithiothreitol (RT, 60 min), alkylated with 54 mM iodoacetamide (dark, RT, 30 min), and digested by adding trypsin at a concentration of 3 ng/μL (overnight at 37 °C), as described in [23].

Peptides were subsequently extracted with 100% acetonitrile (Biosolve), dried by vacuum centrifugation, dissolved in 45 μL 10% formic acid, and spun (10 min, 20000 rpm) prior to analysis. 20 μL of each sample was analyzed on a Q exactive plus (Thermo Fisher Scientific, Bremen) connected to Thermo Scientific EASYnLC 1000 (Thermo Fisher Scientific, Bremen). All columns were packed in-house. The trap column was a double fritted 100 µm inner diameter capillary with a length of 20 mm (Dr Maisch Reprosil C18, 3 μm). The analytical column (Agilent Poroshell EC-C18, 2.7 μm) had an id of 75 µm and a length of 50 cm, and was heated to 40°C. Trapping of the sample was performed at a flow rate of 100 nL/min for 10 min in solvent A (0.1 % formic acid in water), and elution performed with a gradient of 7–38% solvent B (0.1% formic acid 100% acetonitrile) in 75 min, 38–100% B in 3 min, 100% B for 2 min, at a maximum pressure of 800 bar. Nano spray was achieved using a distally coated fused silica emitter (made in-house, o.d. 375 μm; i.d. 20 μm) biased to 1.7kV. The mass spectrometer was operated in the data dependent mode to automatically switch between MS and MS/MS. MS full scan spectra were acquired from m/z 375–1600 after accumulation to a target value of 3x106. Up to ten most intense precursor ions were selected for fragmentation. HCD fragmentation was performed at normalised collision energy of 25% after the accumulation to a target value of 5x104. MS/MS was acquired at a resolution of 17500.

### MS Data Processing

MS raw data were processed using Proteome Discoverer (PD) 2.5.0.400 software (Thermo Scientific, Bremen, Germany). Protein identification analysis was performed using data from UniProt protein sequence database for the Homo sapiens Reviewed Proteome 2024_01 (83,385 entries) along with a common contaminant database from MaxQuant (version 2.6.7.0, Max Planck Institute of Biochemistry, Munich, Germany). FASTA files for the mutant versions of APP were also included to ensure accurate protein identification. Two protein search algorithms were applied: (i) the mass spectrum library search software MSPepSearch, with the NIST human HCD Spectrum Library and (ii) the Sequest HT tandem mass spectrometry peptide database search program. Both search nodes used an ion mass tolerance of 10 ppm for precursor ions and 0.02 Da for fragmented ions. The maximum allowed number of missing cleavage sites was set as 2. Cysteine carbamidomethylation was defined as a constant modification, while methionine oxidation, asparagine and glutamine deamidation, peptide N-terminus cyclization (Gln->pyro-Glut), protein N-terminus acetylation, and methionine loss (Met-loss and Met-loss + Acetyl) were defined as variable modifications. Peptide confidence was set to high, the Inferys rescoring node was applied. Data was validated by the Percolator node.

Protein-label-free quantitation was performed with the Minora feature detector node at the processing step. Precursor ion quantification was performed at the consensus step with the following parameters: peptides = unique plus razor; precursor abundance based on intensity; normalization mode based on the total peptide amount; and pairwise protein ratio calculation and hypothesis testing based on a background-based t-test (38,39).

The dataset of 1,561 proteins identified by the Proteome Discoverer across all conditions (six protein lists: APPWt, APPSA, APPSE, either at day 3 or 7 of differentiation), was further filtered in R Studio (version 4.4.3) to retain only high-confidence identifications. Only master proteins with ‘high’ FDR confidence were further considered. Proteins with only one unique peptide were discarded, unless they presented a molecular weight (MW)≤20 kDa and a sequence coverage of at least 20%, to minimize bias against smaller proteins in peptide-based identification (40,41). Common contaminant proteins and proteins associated to hair, skin/epidermis, tongue and gums (UniProt IDs: P02533, P13645, Q5T749, P35527, Q92764, P04264, Q7Z794, P13646, P19013, Q9NSB4) were removed.

Following, only proteins present in two biological replicates were considered, and nonspecific APP interactors/proteins likely binding to the GFP tag, were further excluded. For this, the mean of each protein’s abundance in APPWT, APPSA and APPSE conditions were calculated, and their ratios to their respective abundances in control (GFP tag alone) were computed. Only proteins with a resulting fold change ≥1.5 relative to control were retained for further analysis.

Proteins were considered ‘specific’ of a condition if only identified in that condition (either APPSA or APPSE); proteins were considered ‘enriched’ in a specific APP mutant if their detected abundance was ≥ 2 relative to the other mutant. Total APP interactome includes 233 proteins identified on this work.

### Bioinformatic analyses

A Principal Component Analysis (PCA) was performed to assess clustering and overall variability among the experimental groups. Further, functional classification of the interactors based on the Protein Class ontology was performed with PANTHER (Protein ANalysis THrough Evolutionary Relationships) database. Significantly enriched Gene Ontology (GO) terms, specifically Biological Processes (BP), were analysed using the ClueGO plugin (version 2.5.10; terms updated on 5^th^ May 2025) in Cytoscape (version 3.10.3) (42,43), (44,45). The proteins lists corresponding to interactors identified at day 3 and 7 of differentiation for each group here considered – APPSA, APPSE and Total APP interactome (all APP interactors pulled down in the three conditions: Wt, SA, SE), were input into ClueGO simultaneously as separate clusters, and a Functional Clustered analysis was performed to enable comparison of functional enrichment between the two timepoints within each group. The following parameters were applied: a minimum of 3 genes per term or 4% of the term’s genes present; merging of redundant terms was activated, meaning terms with more than 50% overlap were fused, leaving only the most significant term based on p-value.

Pathway overrepresentation analyses were performed on g:Profiler (version updated on Jun 16, 2025) (46). For each group and timepoint, the protein lists were input separately, and KEGG, Reactome and WikiPathways were used as pathway sources (Agrawal et al., 2024; Griss et al., 2020; Kanehisa & Goto, 2000; Kolberg et al., 2023). Enrichments were assessed using the hypergeometric test, with p-values adjusted by the Benjamini-Hochberg correction. Only terms with an adjusted p-value ≤ 0.05 were considered statistically significant, using *Homo sapiens* [9606] proteins detected in the brain according to The Human Protein Atlas as background (47).

Protein-protein interaction (PPI) networks were constructed for the interactors of groups ‘Total’, APPSA and APPSE, at each timepoint. PPIs were constructed using *Homo Sapiens* data from BioGRID 5.0 (updated on Dec. 25th, 2025) (48), and analysed in Cytoscape (version 3.10.3) (43). The Cytoscape plug-in clusterMaker2 (49) was used to identify densely connected regions within the PPI networks using the MCODE algorithm, with default parameters (Degree Cutoff = 2, Node Score Cutoff = 0.2, K-Core = 2, Max Depth = 100). For the exclusive and enriched APPSE interactors, proteins were grouped into tables based on common cellular processes and reported roles on neuronal differentiation. The Uniprot database was used to manually retrieve each protein’s name, associated GO-BPs, and reported expression and involvement in disease (50).

The protein summary information was consulted in NCBI for the individual proteins. Phenotype information was obtained from the Mouse Genomics Informatics (MGI) database, which integrates comprehensive genetic, genomic, and phenotypic data from laboratory mouse studies (51). Proteins Q96QA5, P68871, A0A0C4DH42, A0A0B4J1Y9 and P0CG12 were excluded from Table 1, because, although expressed in the brain, some are likely contaminants, while others lack sufficient annotation or evidence linking them to neuronal development.

**Table 1.** Functional clustering of enriched or exclusive APPSE interactors at D3 and D7. The proteins were classified based on evidence in articles and Uniprot linking them to a direct role in the functional groups. *, Proteins with an indirect role in the associated functional group. Diff., differentiation. Ref. References. Cluster 6 information at D3 is in italic as the indicated functions are indirect, derived from the protein’s main functions (induction of translation of specific proteins).

| Functional Group | Dif. day | Proteins | Cellular Event(s) | Main Function | Potential Role in Neuronal Differentiation | Ref. |
| --- | --- | --- | --- | --- | --- | --- |
| **1. Signalling: ligands, receptors, phospho-regulation, transcriptional regulation** | **D3** | ATXN2, DCD, H1-2 | Signal sensing and PTM-mediated modulation; Gene expression control, histone modification. | Translate extracellular and intracellular cues into rapid chromatin and post-translational modifications that change transcription factor activity and mRNA handling. | Regulate neuronal gene accessibility and epigenetic reprogramming; Chromatin reshaping to permit neuron-specific transcription; Influence receptor abundance and trafficking. | (59–63) |
|  | **D7** | ATXN2, FUBP3, NCOA5, PPP1CB | Modulation of Ser/Thr dephosphorylation, regulation of nuclear localization of key transcriptional regulators and chromatin remodelling. | Integrate growth and survival signals with nuclear co-regulator activity to adjust phosphorylation states, modulate chromatin remodelling and transcription factor function, fine-tune promoter and enhancer output. | Control cell-cycle exit and survival programs, stabilize cytoskeletal regulators through phospho-dependent interactions (via PPP1CB), and shape receptor abundance during differentiation, promoting neuronal differentiation signalling. | (63–67) |
| **2. Pre-mRNA processing & spliceosome** | **D3** | DDX1, DDX46, PHF5A, PPIH, PRPF31, RTCB, SF3B5, SF3B6, SNRPB | Pre-mRNA processing, spliceosome assembly. | RNA splicing, snRNP stabilization, RNP remodelling. | Essential for neuron-specific transcript generation and alternative splicing programs during fate commitment; regulate isoform diversity critical for early differentiation. | (68–75) |
|  | **D7** | DHX15, PRPF6, RBM27, RBMX, SF1, SYMPK, U2AF2, WTAP | Spliceosome assembly, splice site recognition, 3'-end/processing scaffold, ribonucleoprotein remodelling. | Ensure precise intron removal and generation of neuron-specific mRNA isoforms by assembling and stabilizing snRNP complexes and coordinating splicing with 3'-end processing. | Drive production of mature, neuron-specific isoforms and alternative splicing programs required for late differentiation and functional maturation. | (76–83) |
| **3. RNA stability, transport (local translation) and translational machinery** | **D3** | ATXN2, ELAVL4, FXR1, FXR2, RPS2, RPS27 | RNA transport and localization to neurites, regulated initiation/elongation of translation, ribosome assembly and direct rRNA/protein methylation. | RNA-binding proteins that regulate mRNA localization, stability, and translation efficiency; Ribosomal structure support translation of synaptic and cytoskeletal proteins during early differentiation. | Maintain localized protein synthesis programs that drive neuronal polarity, growth cone dynamics and early neurite extension. | (84–91) |
|  | **D7** | ATXN2, FUBP3, RBM27, RBMX, RPS12, RPS27 | mRNA stabilization, localization, part of ribonucleoprotein granules, regulated translation; Ribosome structure/function. | RNA-binding proteins that control mRNA availability and local translation in neurites; coordinate translation of transcripts needed for synapse formation. Ribosomal proteins RPS12 and RPS27 contribute to ribosome integrity and efficient translation during maturation. | Maintain transcript pools and enable localized protein synthesis essential for neurite extension and synaptic maturation. Support increased, regulated translation of synaptic and structural proteins. | (87,90,92–96) |
| **4. Proteostasis, PTMs, stress & inflammatory/immune response** | **D3** | DCD, FXR1, PPIH, S100A8, SSBP1 | Protein folding, proteostasis, stress, defence and repair responses. | Maintain proteome integrity via PTMs and peptidyl-prolyl isomerization; Mount stress/inflammatory responses (S100A8, GSDMA, DCD) that impact survival and remodelling. | Support protein biogenesis/folding under differentiation stress and coordinate adaptive inflammatory/stress signalling that can modulate neurite stabilization and survival. | (97–101) |
|  | D7 | GANAB, PRKCSH, SSBP1 | N-glycan trimming and calnexin/calreticulin chaperone cycle; ER quality control for secretory and membrane proteins. | Trim glucose residues from nascent N-linked glycans to permit binding/release cycles with calnexin/calreticulin, promoting correct folding or targeting misfolded glycoproteins for ER quality control. Contributes to ER homeostasis under proteotoxic stress. | Support the secretory pathway during the high demand for synaptic and adhesion molecules and ensure that only properly folded glycoproteins reach the neuronal surface during maturation. | (99,102,103) |
| 5. Mitochondrial function & metabolic support | D3 | FASTKD3, NIPSNAP1, SSBP1 | mtDNA maintenance; mitochondrial RNA processing. | mtDNA replication / mitochondrial RNA handling; mitophagy signalling, energy regulation; quality control. | Provide metabolic support for neurite elongation and axonogenesis; mitochondrial RNA processing adapts energy output during lineage transition. | (99,104,105) |
|  | D7 | SSBP1 | mtDNA maintenance; mitochondrial RNA processing. | mtDNA replication / mitochondrial RNA handling; mitophagy signalling, quality control. | Supply of ATP and Ca<sup>2+</sup> buffering for neurite growth and synaptogenesis; ensure mitochondrial quality during high metabolic demand. | (99,106) |
| 6. Cytoskeleton & membrane remodelling (includes neurite outgrowth and maturation) | D3 | Indirectly: ELAVL4*, FXR1* | Cytoskeletal remodeling, neurite extension. | Coordinate local synthesis of cytoskeletal and adhesion proteins (e.g. GAP43, Tau, CaMKIIa, DSP, ...) to drive neurite extension and synapse formation. | Promote axon/dendrite growth, growth-cone guidance and maturation of synaptic structures. | (101,107–109) |
|  | D7 | INA, PLEKHA6, TUBA1B | Intermediate filament formation, microtubule polymerization, membrane-cytoskeleton scaffolding, cell adhesion. | Structural remodelling, neuriteogenesis, axon/dendrite stabilization, trafficking, and guidance for synaptogenesis. | Directly enable neurite extension, spine formation and membrane trafficking. | (63,110–113) |

APPSE interactors “Q96QA5”, “P68871”, “A0A0C4DH42”, “A0A0B4J1Y9” and “P0CG12” were excluded from Table 1 since some of them are probable contaminants, and others miss detailed information or valid processes concerning neuronal development.

### APP-GFP/interactor subcellular distribution assays and cell morphology studies

SH-SY5Y cells were seeded on poly-D-Lysine (Sigma Aldrich, P6282) precoated coverslips, differentiated, and transfected with APPSE or APP-SA at 2 or 6 DIV, as previously described. At 3 and 7 DIV, cells were fixed with 4% paraformaldehyde for 15–20 min at RT. Samples were further permeabilized with 0.2% Triton X-100 in PBS for 10 min and blocked with 3% bovine serum albumin (BSA) in PBS for 1 h. Primary antibodies diluted in blocking solution (1:50) were incubated for 2 h at RT: mouse anti-HuD (alias ELAVL4, E-1, sc-28299), mouse anti-FXR2 (1G2, sc-32266), mouse anti-α-internexin (G-9, sc-271302), anti-FBP3 (alias for FBUP3, E-8, sc-398466) (Santa Cruz Biotechnology), rabbit anti-ATXN2 (GeneTex, GTX130329), rabbit anti-α-tubulin (TU-01, Novus Biologicals, NB500-333). Secondary Alexa Fluor 594-conjugated goat anti-mouse (A11005) and anti-rabbit (A11012) antibodies (Invitrogen), diluted in blocking solution (1:300), were incubated for 1 h at RT. Coverslips were mounted with DAPI-containing Vectashield antifading mounting medium (Vector, H-1200) and images were acquired using a Zeiss LSM 880 Airyscan confocal microscope (100x oil objective).

Morphometric analysis was performed on differentiated SH-SY5Y cells, transfected with the three APP-GFP cDNAs (Wt, SE or SA) at 6 DIV and fixed at 8 DIV, for neurites to have more time to develop under a specific APP overexpression background. 30 randomly selected digitized images were analyzed per sample, with an average of 40 transfected cells (GFP expressing) per biological replica of each condition (Wt, SE, or SA). Measurements were performed using ImageJ XXX on matching PhC microphotographs. The number and length of processes per cell were quantified and categorized as follows: <20 µm, 20-35 µm, 35-50 µm, and ≥50 µm. Processes ≥35μm were already considered neurites. Processes mean length only includes processes ≥20 µm.

## Results

### S655- and differentiation time-dependent APP interactors and their functional classes

This study aimed to gain insight on the role of APP S655 phosphorylation in neuronal differentiation, by identifying APP protein interactors dependent on its S655 phosphorylation state. Neuronal-like differentiated SH-SY5Y cells, overexpressing Wt or S655 phosphomutants APP-GFP cDNAs for 24h prior to collection, were used as the protein pool. The GFP Trap assay was used to immunoprecipitate the APP proteins (APPWt, APPSA, and APPSE) and their respective interacting partners, which were subsequently identified by mass spectrometry.

In total, 233 proteins were identified across all groups and timepoints, of which 64 were already described APP interactors, while 169 represent newly identified APP interactors. The number of interactors identified for each group and timepoint, along with their intersections, are shown in Venn diagrams (Figure 1A). From differentiation day 3 (D3) to day 7 (D7), APPWt and APPSE groups present a relatively stable number of interactors (n=110→108 and n=78→72, respectively), whereas the number of APPSA interactors increased substantially (n=97→163). Of note, only proteins appearing in at least two replicates were here considered; if all interactors with a fold change >1.5 relative to the GFP control were considered, all groups would exhibit an increase in interactor number over the course of differentiation (Supplementary Figure 1A), suggesting that the amount of APP interactors increases as the cell specializes into a neuron. The number of APPWt interactors is intermediate to the ones of the phosphomutant groups, consistent with the fact that APPWt can be either S655 phosphorylated or not (Figure 1A). As such, both APPSE and APPSA share most of their interactors with APPWt on D3: 83.3% for APPSE and 86.6% for APPSA (with 60 interactors common to the two phosphomutants). This overlap decreases to 50.3% (APPSE) and 58.3% (APPSA) at D7, suggesting that S655 phosphorylation imposes more specification as the cells become more neuronal-like, negatively regulating a higher number of APP interactions at D7. Consistent with a specialization of the APP interactome upon S655 phosphorylation, Principal Component Analysis (PCA) of all replicates showed APPSE samples tightly clustered, APPSA samples widely dispersed, and APPWT samples clustered either with APPSE or with APPSA (Supplementary Figure 1A). The main proteins contributing for groups separation at each day are presented in Supplementary Figure 1B. APPSE maintained ∼20% exclusive interactors at both D3 (18, 23.1%) and D7 (16, 22.2%), while the number of APPSA exclusive interactors was higher and markedly increased from D3 (37, 38.1%) to D7 (107, 65.6%). Figure 1B shows that the identity of APP interactors partially changes as the cells acquires neuronal traits, such as a more neuronal-like proteome and morphology. For all the subsequent analyses, the identified interactors were subgrouped into three categories: 1) Total APP interactors, which includes all APP interactors identified in this work (131 at D3; 199 at D7); 2) APPSA interactors (97 at D3; 163 at D7); and 3) APPSE interactors (78 at D3; 72 at D7).

**Figure 1.**
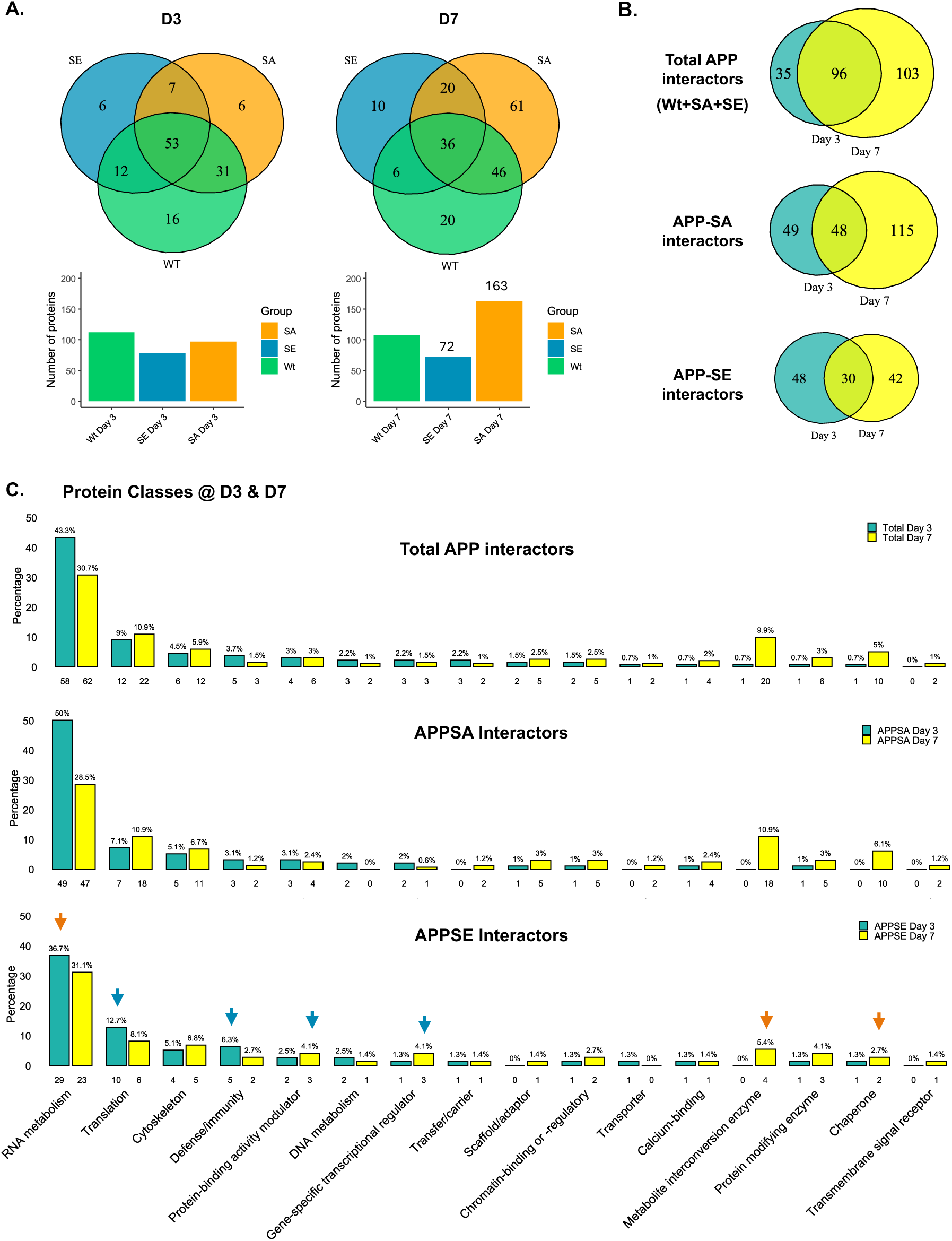
APP protein interactors change with S655 phosphorylation state and differentiation time. **A.** Venn diagrams presenting the numbers of interactors experimentally detected in the pull-downs of APPWt and the S655 phosphomimetic mutants (APPSA and APPSE), and the intersections between these conditions, at both differentiation days. Bellow: Barplots representing the number of interactors in the different groups on each differentiation day. **B.** Comparison of the numbers of APP interactors (APP included) on each differentiation day for Total, APPSA and APPSE interactors. **C.** Protein class classification according to PANTHER (Protein ANalysis THrough Evolutionary Relationships) for ‘Total’, APPSA and APPSE groups. D3 and D7, days 3 and 7 of neuronal differentiation. Only classes with n=2 interactors at t least one of the differentiation days were included.

To start attributing putative functions to the phosphomutant interactors, a functional classification was performed based on the “Protein Class” ontology from PANTHER (Protein ANalysis THrough Evolutionary Relationships) database (Figure 1C). In the Total APP interactors group, the dominance of RNA metabolism proteins is notable at both days, with around 60 interactors related to this class in each time point. Other classes mainly represented across differentiation time are Translational (9% at D3; 11% at D7) and Cytoskeletal proteins (4.5% at D3; 5.9% at D7). At D3 only two other classes have ≥3% of interactors, namely Defense/immunity (3.7%) and Protein-binding activity modulator (3%). Some classes increased from D3 to D7, particularly ones related to post-translational modifications (PTMs), such as Metabolite interconversion enzymes (from 0.7% to 9.9%) and Protein modifying enzymes (0.7% to 5%), followed by Chaperones (0.7% to 3%). APPSA interactors account for most of the total APP interactors, which explains why their protein class distribution is very similar to the Total APP interactors one, with RNA metabolism proteins being the most abundant, followed by Translational and Cytoskeletal proteins, at both differentiation time points. Apart from a more pronounced decrease in RNA metabolism with time of differentiation, the remaining distributions for APPSA interactors mirror the total APP profile, with a similar increase in PTM-associated proteins (including metabolism-related ones). The APPSE group, however, presented some particularities. Although RNA metabolism remained the dominant class, it only represented one-third of the classes (orange arrow) and this class only slightly decreased at D7 (from 36.7% to 31.1%). At D3, proteins related to Translation (12.7%; vs 7.1% for APPSA) and Defence/immunity (double % relatively to APPSA) were more abundant in this group (blue arrows). Of note, the Defence/immunity class is mainly composed of ribosomal protein subunits (RPS) and one chaperone, strengthening a pS655-dependent enrichment of Translation-related functions. At D7, there was an APPSE-specific growth in the Gene-specific transcriptional regulator (from 1.3% to 4.1%) and Protein-binding activity modulator (from 2.5% to 4.1%) classes, a trend not observed in APPSA. Conversely, the Metabolite interconversion and Chaperones classes increased less in percentage than in APPSA.

### Biological Processes and Pathways of APP S655 phosphomutants protein interactors

Subsequent functional enrichment analyses were conducted to explore the biological processes (BP) and pathways enriched in the APP protein interactors groups. Figure 2 shows the APPSA-and the APPSE-enriched BP terms, colour-categorized by differentiation day. Terms were classified as mainly enriched in D3 (not found) or D7 (yellow nodes) if ≥75% of their protein members were detected only on that day. Otherwise, terms were considered enriched in both days (green nodes).

**Figure 2.**
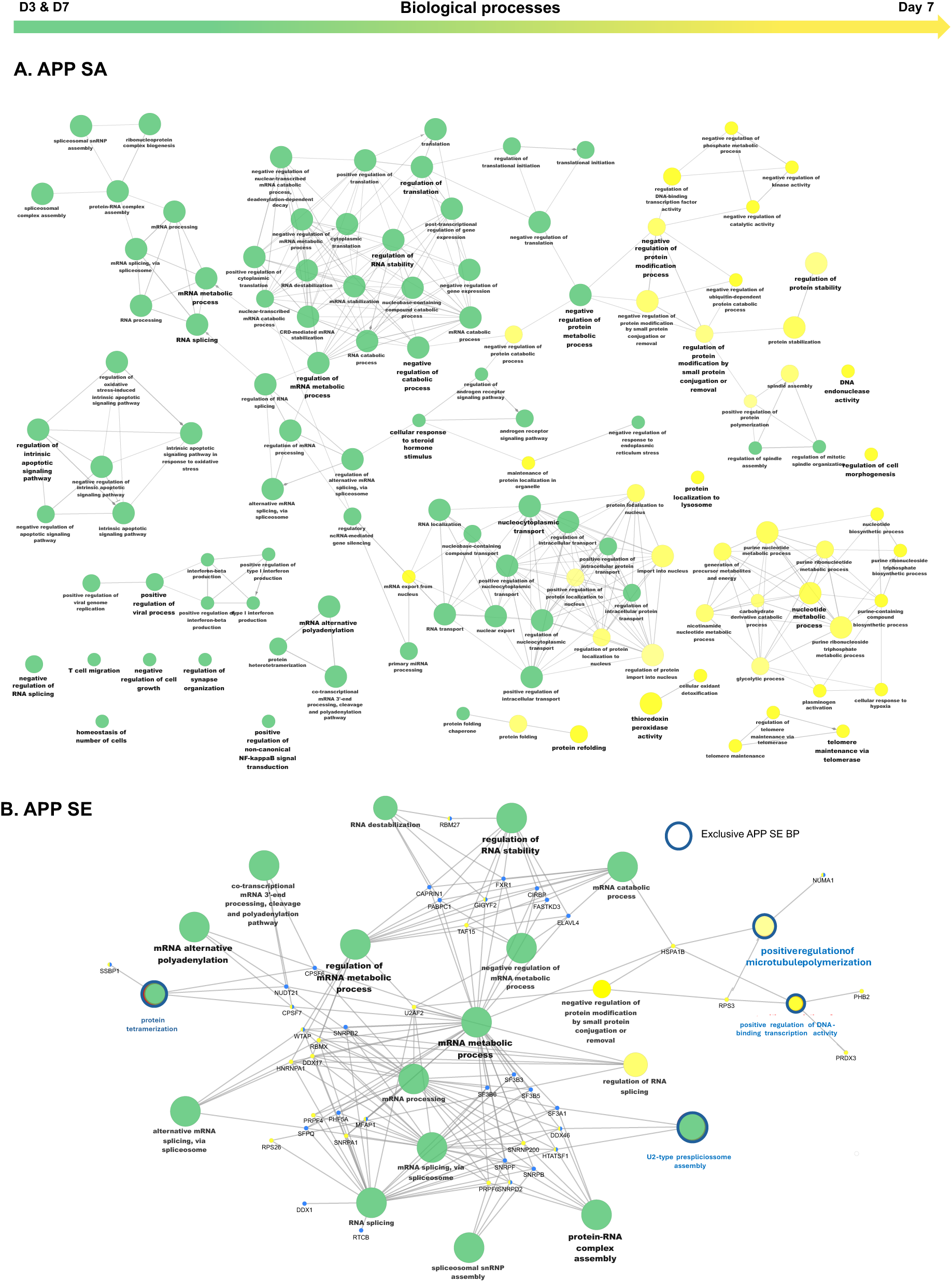
Biological processes (BP) gene ontology (GO) terms for the APPSA (A) and APPSE (B) total groups. BPs specifically enriched at both differentiating days (D3 & D7) are presented in green and those enriched at D7 are in yellow. No terms only enriched at D3 were found. Terms were considered enriched in only one day if 75% or more of its protein members were found only in that day. Only terms with p<0.05 upon Benjamini-Hochberg multiple test correction are represented. The size of the term nodes is based on term’s significance (higher, the lower its p-value). Terms with more than 50% overlap are fused and only the more significant term is presented in this network.

The APPSA BP network was composed of 120 terms, 86% of which exclusive (not detected in APPSE). Prominent in this network were processes related to RNA metabolism, stability, transport and polyadenylation, together with clusters associated with translational initiation and nuclear-cytoplasmic transport/localization (all shared between D3 & D7; the nuclear-cytoplasmic transport/localization become more specialized at D7, with added members). Processes mainly enriched at D7 were related to PTM (mainly negative regulation of phosphorylation), protein stability and nucleotide metabolism. There were also smaller clusters of processes related to apoptotic signalling regulation, cellular response to steroid hormone, innate defence (all these shared at D3 & D7), plus protein refolding and telomere maintenance (enriched at D7). Relevant isolated BP terms of APPSA interactors included “Regulation of synapse formation”, “Positive regulation of NF-kB signal transduction” (both days), plus “Protein localization to lysosome” and “regulation of cell morphogenesis” at D7.

The network of enriched BP associated to the APPSE interactors was formed by 20 terms, most of them shared with APPSA (80%, non-delineated circles). Like APPSA, APPSE interactors were strongly associated with RNA-related processes on both days, particularly mRNA metabolism, splicing and stability, and protein–RNA complex assembly. Processes enriched at both days but exclusive of APPSE were “Protein tetramerization” (that shared proteins with an RNA processing cluster) and “U2-type prespliceossome assembly” (sharing proteins with a spliceosome-related cluster). A role in RNA splicing was further strengthened with the D7-exclusive term “Regulation of RNA splicing”. Interestingly, two other terms were exclusive to phosphoS655 at D7: “Positive regulation of microtubule polymerization” and “Positive regulation of RNA-binding transcription factor activity”.

Pathway enrichment analysis retrieved 47 pathways for APPSA interactors and 56 for APPSE interactors at D3, and 113 (APPSA) and 21 (APPSE) pathways for D7 interactors. Many of these were shared by both groups, with the top 20 shared pathways all related to RNA processing and translation, at both differentiation days (Supplementary Figure S2). Nevertheless, at D7 APPSE had a higher percentage of pathways related to translation (APPSE D3: 35.7%, D7: 47.6% vs. APPSA D3: 31.9%, D7: 18.6%) and to RNA metabolism (APPSE D3: 32.1%, D7: 33.3% vs. APPSA D3: 34%, D7: 17.7%). In contrast, at D7, APPSA showed greater representation of pathways related to transcription factors (APPSA D3: 4.3%, D7: 4.4% vs. APPSE D3: 3.6%, D7: 0.0%) and signalling (APPSA D3: 6.4%, D7: 12.4% vs. APPSE D3: 1.8%, D7: 0.0%). Exclusively enriched pathways were observed mainly at D3 for APPSE and at D7 for APPSA. At D3, APPSA had only three exclusive pathways (6.4%), two related to signalling and one to trafficking, whereas APPSE presented 12 exclusive pathways (21.4%). These included eight related to rRNA and translation, two linked to viral host response (potentially translation, given shared proteins with the previous terms), and two associated with neuronal differentiation: “Axon Guidance” and “Nervous system development”, associated to proteins PABPC1, RPS5, RPL38, RPS2, RPS18, RPL13, RPL30 and RPS27 (Supplementary Figure S2, D3). At D7 of differentiation, a stage marked by a boost in APP interactors binding to APPSA, the pattern was reversed and APPSA presented a very high number of pathways, including 93 exclusive ones (82.3%). Top 20 APPSA pathways (prioritized by p-value) were all related to translation, including quality control of mRNA (Supplementary Figure S2, D7). APPSA exclusive pathways at D7 were generally related to translation (15), signalling events (14), RNA metabolism (13), regulation of expression (5), protein trafficking (5), PTM (4) and neuronal processes (4).

### APPSE binding partners form highly interconnected protein-protein interaction networks

Protein-protein interactions (PPI) networks were constructed with all APPSE interactors identified at D3 or D7 (Figure 3A left and right, respectively). Previously described APP interactors are outlined in red, including 9 proteins at D3 and 10 at D7, while the remaining proteins represent novel APP interactors identified in this study (D3: 88.6%, D7: 86.1%). A closer look to the APPSE networks of Figure 3A reveals that six of the known APP binders – DDX1, H1-2, S100A8, SF3B6, and FUBP3 (52,53)– are exclusive or enriched APPSE binders, with the first two having FXR1 as a common binder (54–56). FXR1 also binds the D3+D7 exclusive SE interactor ATXN2, to which FUBP3 also binds (Figure 3A and B) (54).

**Figure 3.**
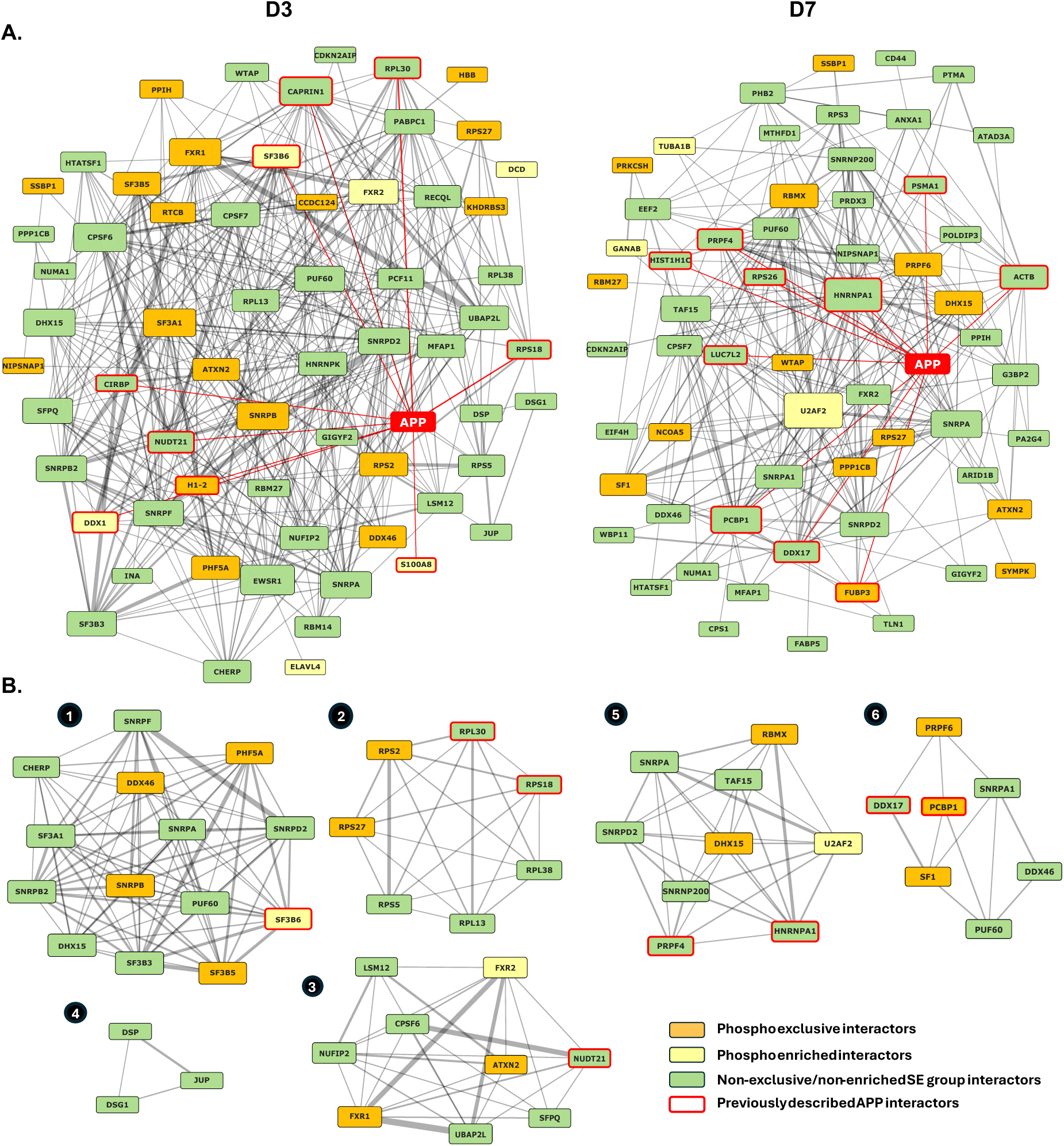
APPSE protein-protein interaction (PPI) networks at D3 and D7 of neuronal differentiation. **A.** Full APPSE interactors PPI network. **B.** Clusters formed by MCODE using the full APPSE networks. Node sizes in **A.** and **B.** are based on the node degree of each interactor in the total APPSE network.

The D3 APPSE PPI network comprised 64 nodes connected by 340 edges, with a clustering coefficient of 0.42, indicative of a moderately modular organisation with a large interconnected component and a few peripheral groups. The D7 network had 60 nodes and 245 edges and, although the average number of neighbours slightly decreased from 10.6 to 8, the clustering coefficient remained the same (0.43), as well as the path length (2.2 for both days). Both networks connected most of the APPSE interactors identified in this work, revealing that many of these had already been reported to interact and likely participate in common functions. Only a small part of APPSE binders, 15 at D3 and 2 at D7, were excluded from these PPI visualizations, given the absence of known interactors within the PPI. Automatic clustering of Figure 3A networks identified four protein clusters at D3 (Figure 3B, 1-4) and two clusters at D7 (Figure 3B, 5-6). Panther protein class analysis of each cluster indicated that D3’s cluster 1 is composed by RNA metabolism proteins (12 out of 14), cluster 2 consisted of only translation-related proteins, cluster 3 of both RNA metabolism and translation proteins (4 and 2 out of 9, respectively), while the smaller cluster 4 has one cytoskeletal protein (DSP) and a cell adhesion molecule (DSG1). At D7, cluster 6 is mostly composed of RNA metabolism proteins (8 out of 10) and a scaffold/adaptor protein, and cluster 2 is also mainly composed by RNA metabolism proteins (6 out of 7) and a gene specific transcriptional regulator. Nearly all these clusters included exclusive and enriched APPSE interactors, indicating that S655 phosphorylation modulates interactions within key APP functional complexes.

To further test if S655 phosphorylation promotes the formation of specific protein-protein complexes, PPI networks with only APPSE exclusive and enriched interactors were subsequently created (Figure 4A). A high number of interactions was visible, with the D3 network presenting 19 interactors (out of 26) and 34 edges, and the D7 network presenting 14 interactors (out of 21) and 19 edges. Hence, at both differentiation days, and using only experimentally validated interactions, more than two thirds of the APPSE exclusive/enriched interactors bind together in a network that includes APP, independently of the common APPSA/APPSE binders (in green in Figure 3).

**Figure 4.**
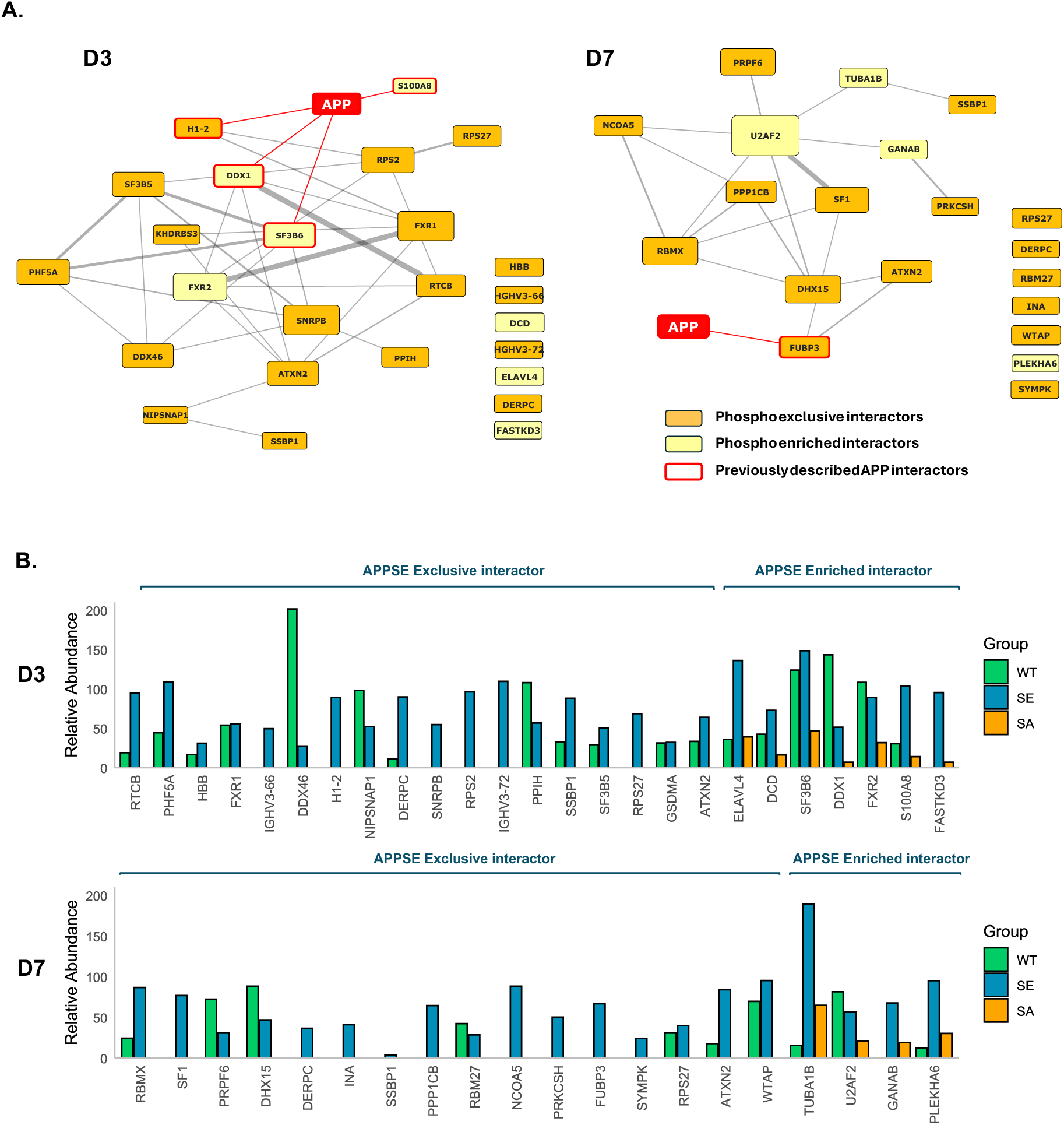
APPSE exclusive and enriched interactors at D3 and D7 of neuronal differentiation. **A.** APPSE exclusive (only detected on APP SE) and enriched (ratio APPSE / APPSA ≥ 2) interactors PPI network; node size in is static. **B.** Relative abundances of exclusive and enriched APPSE interactors in each group (APPWt, SE, SA), at day 3 (D3) and day 7 (D7) of differentiation. Relative abundances are only shown if the interactor is detected on the group (FC≥1,5 relative to control). Average values of a protein’s abundance in the replicate of a group, minus the median abundance of that protein in the empty GFP vector, in same day (D3 or D7).

Figure 4B shows that the abundance of these APPSE exclusive and enriched interactors varied across differentiation time points. At D3, ELAVL4 and SF3B6 presented the highest relative abundances, whereas TUBA1B was the most abundant at D7. Among the APPSE exclusive interactors, 15 proteins (IGHV3-66, H1-2, SNRPB, RPS2, IGHV3-72, RPS27, SF1, DERPC, INA, SSBP1, PPP1CB, NCOA5, PRKCSH, FUBP3, SYMPK) were only co-immunoprecipitated with APPSE and not with APPWt, suggesting that a sustained S655 phosphorylation may be required for their binding to APP. Three of these interactors (DERPC, RPS27, SSBP1), together with ATXN2, were detected as exclusive APPSE interactors at both differentiation days. While the neuritogenesis-associated ATXN2 was the only interactor whose relative abundance increased with differentiation time (from ∼60 at D3 to ∼90 at D7), the abundance of the ribosome biogenesis and mitochondrial stability-associated DERPC, RSP27 and SSBP1 proteins decreased from D3 to D7 (about half for RPS27; SSBP1 became nearly undetectable).

### S655 phosphorylation potential functions in neuronal differentiation

The smaller APPSE interactome suggests a functional specification for this phosphorylation. Further, most of the exclusive and enriched APPSE interactors interact with one another (Figure 4A), suggesting that they may act synergically as part of functional modules relevant to differentiation. To explore this potential functional specification in neuronal differentiation, we performed literature mining to extract the known functions of the APPSE exclusive and enriched interactors, followed by manual functional clustering. Table 1 groups these interactors by cellular process and outlines the proposed roles of each functional cluster in neuronal differentiation.

Spliceosomal and core pre-mRNA processing factors (cluster 2) remained largely present at both time points, as well as proteins annotated for mRNA stability, transport and ribosome/translation functions (cluster 3, with several RBPs and ribosomal proteins). These are related to neuronal differentiation by driving neuron-specific alternative splicing and mRNA processing programs that generate isoforms required for neuronal fate specification, axon guidance and synapse formation. In parallel, RNA-binding proteins, mRNA transport factors and ribosomal components support the localization and activity-dependent local translation of transcripts in neurites, growth cones and presynaptic compartments, all essential for neuronal maturation (57,58).

Although the functional clusters remained the same, there were some fluctuations between D3 and D7, with increases in the representation of signal transducers and nuclear co-regulators (cluster 1) that were accompanied by decreases in the number of proteins in cluster 4 (proteostasis and stress) and cluster 5 (energy, mitochondrial biogenesis and maintenance). Additionally, cytoskeletal and extracellular-matrix/adhesion proteins were only present at D7, since at D3 these functions were only indirectly represented by ELAVL4 and FXR1, that promote translation of mRNAs whose protein products are related to cytoskeleton-remodelling and neuritogenesis.

### PhosphoS655 positively modulates APP-induced neuritogenesis

A graphical representation of the APPSE exclusive and enriched interactors’ potential location and functions on neuronal differentiation at D3 and D7, is presented in Figure 5A. We also performed immunocytochemistry analyses of the potential subcellular regions of APPSE binding to some of these interactors, selected based on their abundance and/or functional relevance (Figure 5B).

**Figure 5.**
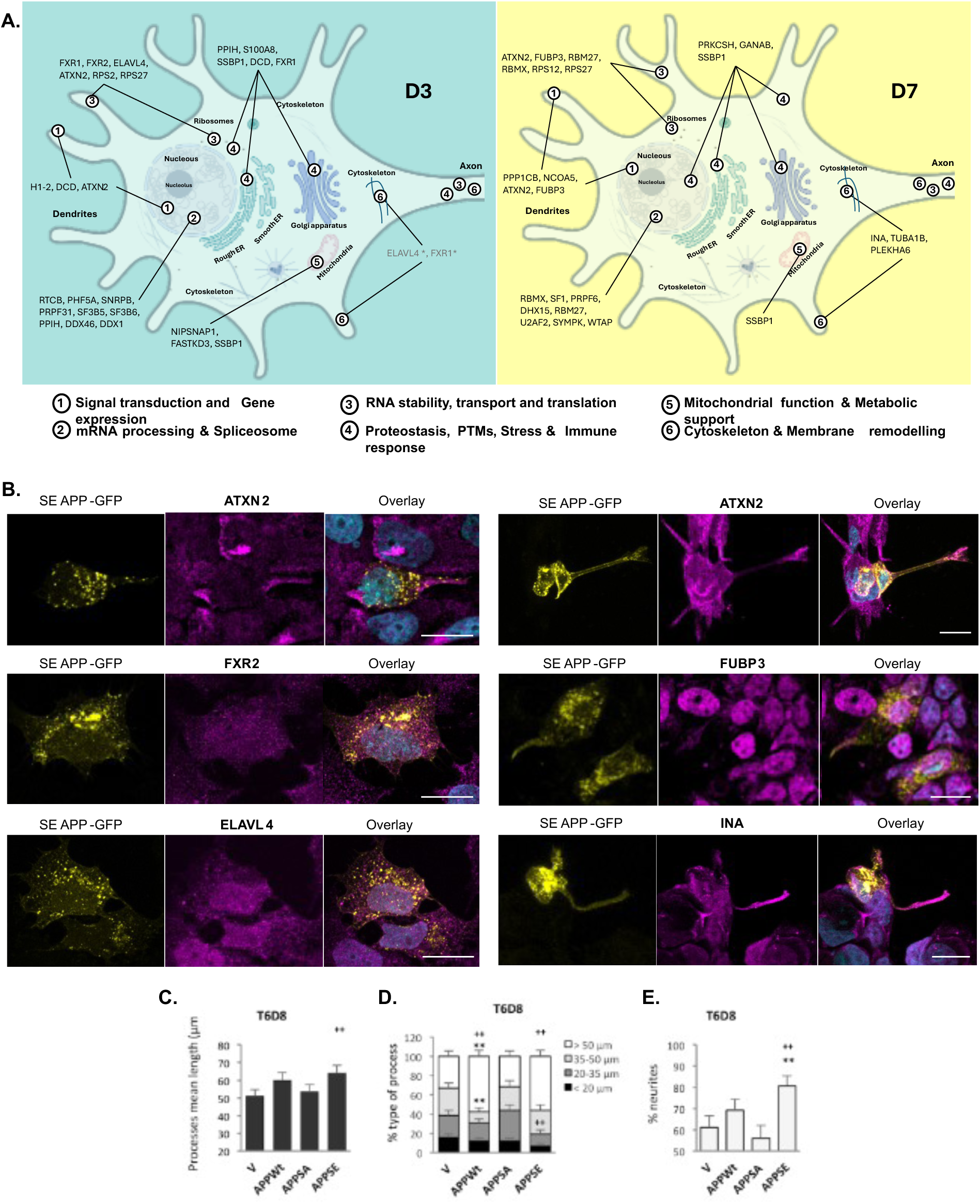
APPS exclusive and enriched interactors organized in functional clusters in neuronal differentiating cells. **A.** Schematic figure created using BioRender.com. The terms listed are: (1) Signal transduction (including ligand binding, receptor trafficking and phosphoregulation) and gene expression, (2) mRNA processing and spliceosome, (3) RNA stability, transport and translation (including local translation and translational machinery), (4) Proteostasis, PTMs, stress & immune response, (5) Mitochondrial function & metabolic support and (6) Cytoskeleton & membrane remodeling (including regulation of neurite outgrowth and synaptogenesis). **B.** Immunocytochemistry analysis of the subcellular distribution of APPSE (GFP tagged) and various of its exclusive or enriched interactors (in magenta) in SH-SY5Y cells differentiated with retinoic acid for 3 (D3) or 7 (D7) days.

Microphotographs of differentiating SH-SY5Y cells at D3 (Figure 5B, left panels) showed **ATXN2** localized in puncta/spots throughout the cell. Its signal was generally faint, except for some regions near the plasma membrane (PM) along existing projections, and at sites from which new projections could emerge, where it appears to co-localize with APPSE. **FRX2**, an enriched APPSE interactor and ATXN2-binder, was localized to the nucleus, cytoplasmic puncta and PM. FXR2 showed some co-localization with APPSE at all these locations, but particularly in perinuclear vesicles and Golgi, as well as in various puncta at the PM. Co-localization was much less evident with APPSA (Supplementary Figure S4). The APPSE-enriched interactor **ELAVL4/HuD**, highly abundant in neurons, displayed a granular cytoplasmic distribution and was highly enriched in the nucleus. ELAVL4 showed some co-localization with APPSE in various cellular puncta, but mainly at perinuclear vesicles/Golgi and PM (including several projections/pre-neurites).

Conversely, ELAVL4 co-localization with APPSA was minimal, and ELAVL4 appeared to be less visible at the nuclei and PM of APPSA-transfected cells (Supplementary Figure S4).

At D7 (Figure 5B, right panels), the APPSE interactors analysed were ATXN2, FUBP3 (alias FBP3, also an ATXN2-binder), and the cytoskeleton-related INA. **ATXN2** displayed a punctate distribution throughout the cell but was highly enriched near the PM, particularly in cell projections/pre-neurites and growth cones. ATXN2 co-localized with both APPSE and APPSA in all these sites (ATXN2 D7 panels in Figure 5B and Supplementary Figure S4), what suggests that ATXN2 does not bind to APPSA due to a lack of affinity, but may participate in some related protein complexes/functions. **FUBP3** was predominantly located in the nucleus (nucleoplasm) and at/near the PM, particularly along projections/pre-neurites and growth cones. FUBP3-APPSE co-localizing spots were visible at the PM and projections, and to a lesser extent in the nuclei. A similar co-localization pattern was observed for FUBP3 and APPSA (Supplementary Figure S4). Finally, APPSE was also observed to co-localize with cytoskeleton-associated proteins like INA (intermediate filament protein with reported interactions with APP (114), despite not listed on BioGrid), in cytoplasm and PM, particularly in membranar projections/pre-neurites. In APPSA-expressing cells, INA staining appeared as small puncta in the nuclei (highly enriched) and cytoplasm, with little to no co-localization with APPSA.

Noticeably, the signal intensity of some of these proteins (e.g. ELAVL4 and INA), appeared to increase in cells with higher levels of APPSE expression, particularly in APPSE ones, what may be due to induced expression and/or reduced degradation, but warrants confirmation but other techniques. Further and related, there were cells where highly expressed APPSE seemed to trap its interactors at the Golgi and cytoplasm (Supplementary Figure 5A). Other important note was that at D7 of RA-induced differentiation, more than half of the APPSA-expressing cells exhibited abnormal morphology (e.g. rounded shape, altered nuclei structure), suggestive of an early apoptotic state. This phenotype was more evident for cells with higher transfection levels, while cells with low to moderate expression seemed less affected.

Given the potential involvement of some of its exclusive and enriched interactors in neuritogenic processes, and some visual indication of a more elongated phenotype in APPSE transfected cells (relatively to APPSA-transfected ones), we assessed the impact of S655 phosphorylation on neurite outgrowth and elongation. The number of protruding processes was scored, and their lengths measured, in APPSE, APPSA and APPWt-expressing cells (Figure 5C-E). APPSE expression favoured the elongation of processes, resulting in a significant increase in their mean length (Figure 5C) and a higher percentage of longer processes (35-50 µm and >50 µm; Figure 5D), an indirect indicator of differentiated neurite-bearing cells (Figure 5E). Notably, overexpression of APPWt already enhanced some of these parameters, while APPSA remained similar to control cells (transfected with empty vector). Conversely, the total number of processes per cell were slightly higher for APPSA cells, although not reaching significance in these conditions.

## Discussion

In this work, we aimed to characterize the S655 phosphorylation-dependent APP interactome under a neuronal differentiation context, using mass spectrometry to identify proteins interacting with different APP-GFP fusion proteins carrying S655 mutations. From the 234 potential APP interactors retrieved by our working methodology, 65 were already known APP interactors (27.7% of ID proteins), including APP itself, which was detected in all groups. Another known APP binder, the PP1 catalytic subunit PPP1CB, was only retrieved in the APPSE group on D7, in accordance with its reported function in dephosphorylating APP at this residue (115). These support that the experimental methodology was sufficiently robust. The results showed a smaller set of APPSE interactors compared to APPSA, indicating that S655 phosphorylation drives functional sub-specialization and/or enrichment in specific functional subsets, while likely preventing some interactions. Differential interactions should be derived from altered conformation of the C-terminus due to the presence of the negatively charged phosphate group within the YTSI sorting motif (116,117), what increases/reduces the affinity of binders, and/or by differential localization of the APPSA vs APPSE species. Our previous studies already showed enhanced trafficking of APPSE from the TGN to the PM and vice versa, while APPSA remained for longer at the Golgi and lysosomes (34,37). Since the APPSE species is more exported to the PM and vesicles (exocytic and early endosomes), it is expected to interact less with proteins abundant in the ER and Golgi apparatus, for example. Regarding the nature of APP interactors, functional enrichment analyses revealed that the ‘Total interactors’ and the APPSA groups were very similar, in line with expected temporary occurrence of phosphoS655 APP. In all groups and timepoints, the main functions of APP interactors were related to RNA metabolism (including splicing) and translation (including ribosomal structure). At D7, the APPSA interactome was associated to signalling mechanisms (including apoptotic) and protein localization terms, including the “Protein localization to lysosome” consistent with the expected prolonged localization of APPSA in this organelle (34). Overall, APPSE seems to be more associated to specific translation events, microtubule and neuronal terms.

Nuclear functions related to RNA processing are likely associated with the AICD fragment rather than the full-length APP, since it is the main form trafficked to the nucleus. AICD has been implicated in nuclear transcriptional regulation through the formation of a complex with the adaptor protein Fe65 and the histone acetyltransferase Tip60 (118), although its precise contribution for transcriptional regulation remains debated (119). Colocalization studies have shown that APP CTFs, such as AICD, accumulate within intranuclear compartments enriched in splicing factors, suggesting a potential involvement in alternative splicing regulation (120). Given that the human brain exhibits a uniquely high level of alternative splicing, changes in splicing decisions can significantly influence protein structure, mRNA localization, translational efficiency, and decay. In neurons, such regulatory networks are essential for their development and for maintaining polarity and synaptic plasticity. APP may interact with diverse RNA-binding proteins (RBPs) or be incorporated into messenger ribonucleoprotein complexes to affect the post-transcriptional fate of multiple transcripts (119,121). The APPSE interactome was observed to be enriched in pre-mRNA processing and spliceosomal components (e.g. SF3B5/6, PRPF6, U2AF2, DHX15, PPIH), suggesting that the AICD fraction generated from APPSE may preferentially engage with these nuclear splicing hubs(124).

Regarding the APPSE-specific or enriched partners, these were observed to form highly interconnected, dense PPI clusters, and many have known functions in neurite outgrowth. Some co-localized with APPSE not only at perinuclear vesicles, but also at the PM, growth cones and (pre)neurites. Additionally, APPSE expressing cells generally have pre-neurites and neurites more elongated. All this indicates that S655 phosphorylation likely induces the formation of specific APP-including “neurodifferentiation hub” (protein macrocomplexes with role in neuritic elongation) rather than only simply adding or removing single binders to APPSA-containing complexes/functions.

A central APPSE-exclusive interactor at both time points is ATXN2, a multifunctional RBP involved in RNA metabolism, stress granule dynamics, endocytosis and calcium-related signalling (122). ATXN2 binds factors such as EGFR, known for its neuritogenic properties, ACTN1, DDX1, FXR1/FXR2, and FMRP, linking APPSE to growth-factor signalling, actin remodelling, endocytosis, and stress-responsive RNP condensates that regulate mRNA stability and translation in neurites (8,54,86,123). Given ATXN2’s known roles in stabilizing specific mRNAs and organizing stress granules, its selective association with APPSE suggests that pS655 APP is preferentially recruited into RNP assemblies that coordinate receptor signalling (e.g., EGFR), actin dynamics and RNA handling during early neuronal differentiation. This is consistent with our imaging data, where ATXN2 puncta accumulate near membranes, growth cones and actin-rich structures, co-localizing with APPSE in both early projections and more elaborated neurites.

A particularly striking feature of the APPSE-enriched interactome is the predominance of RBPs and translational regulators with well-established roles in neurite outgrowth and synaptic development. ELAVL4/HuD, FXR1, FXR2 and FUBP3, all appear as SE-exclusive or enriched interactors at different time points. ELAVL4/HuD stabilizes and promotes the translation of key neuronal mRNAs, including GAP-43, Tau and NRN1, which are required for growth cone formation, axon extension and regeneration. Decreased HuD function impairs GAP-43 mRNA stability and leads to defective neurite outgrowth, while HuD upregulation is associated with enhanced neurite extension, recovery after injury, and decreased Aβ production (107,125). Concordantly, phosphorylation of APP at S655 has been linked to decreased Aβ production (37,126). Notably, within the D3 APPSE interactor network is EWSR1, a transcription repressor that interacts with ELAVL4/HuD. Disruption of EWSR1 function in knockout mouse and zebrafish models results in severe developmental phenotypes, underscoring the importance of this interactor for neuronal development and survival (127,128). The preferential association of ELAVL4 with APPSE at D3, and their co-localization at projections and PM suggest that pS655 APP may scaffold a HuD-containing complex that boosts local expression of growth cone and cytoskeleton regulators when neurites start to emerge.

The Fragile X family proteins FXR1 and FXR2 are involved in translational control at neurites. FXR1 and FXR2 regulate mRNA translation efficiency, often through miRNA-dependent mechanisms, and share targets and structural features with FMRP. This protein, for instance, controls Map1b mRNA translation during Sema3A-mediated axon guidance and interacts with APP and CaMKII pathways via CYFIP1/eIF4E complexes (129). FXR1 displays a distinct expression profile, being more prominent in muscle and certain neuronal populations, and has been implicated in translation regulation, RNA stability and local mRNA control in response to signalling cues (108). FUBP3 binds 3′ UTR elements to modulate translation of specific transcripts, has been linked to neuronal functions (e.g. FGF9) and, more recently, to amyloid-β related signalling in neurons (92,130).

At later differentiation stages (D7), more APPSE interactors are associated to cytoskeleton and potentially to membrane/ECM coupling, with INA, TUBA1B and PLEKHA6 appearing as exclusive/enriched partners. INA (a neurofilament protein) and TUBA1B (αtubulin) are structural elements required for neurite elongation and vesicular transport, with alterations in their normal physiology potentially leading to neurodegeneration (110,131,132).

In conclusion, phosphorylation at S655 changes APP binding partners over the course of neuronal differentiation, acting like a switch to increase or decrease APP affinity to specific protein complexes. These phosphorylation-driven changes in protein binding suggest that APP can take on different functional roles at different stages of neuron development, affecting specific RNA processing and protein production in the cell, with consequences for neuritogenesis, namely APPSE-driven neurite elongation. While these findings are preliminary and mainly hypothesis-generating, they provide testable directions for follow-up experiments to understand how S655 phosphorylation controls APP and cell behaviour.

## Acknowledgments

This work was supported by the PT Foundation for Science and Technology (FCT, I.P.), and the European Regional Development Fund (FEDER), via programs Portugal2020 and 2030, Centro2020 and 2030 and COMPETE2030, by funding the Institute of Biomedicine (iBiMED; UIDB/04501/2020) and projects GoBack (PTDC/CVT-CVT/32261/2017) and Reconnect (COMPETE2030-FEDER-00891600). The MS experiments were performed in the Biomolecular Mass Spectrometry and Proteomics group at Utrecht University (NL), under a grant of the PRIME-XS consortium (PRIME-XS 292). Microphotographs were acquired in the LiM Platform for Advanced Optical Imaging of iBiMED-UA, member of the Portuguese Platform for Bioimaging (PPBI), a node of the EuBi European Bioimaging Network.

