## Supplementary Figures for "The phosphoS655 Alzheimer’s Amyloid Precursor Protein (APP) interactome in neuronal differentiation"

**A.**

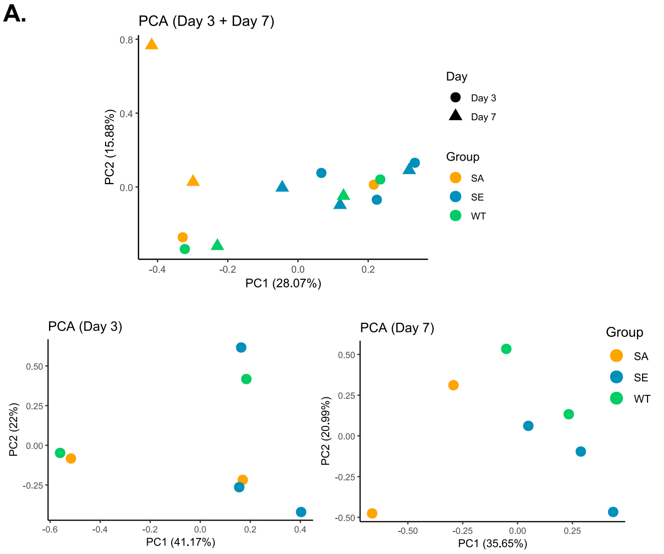


**C.**

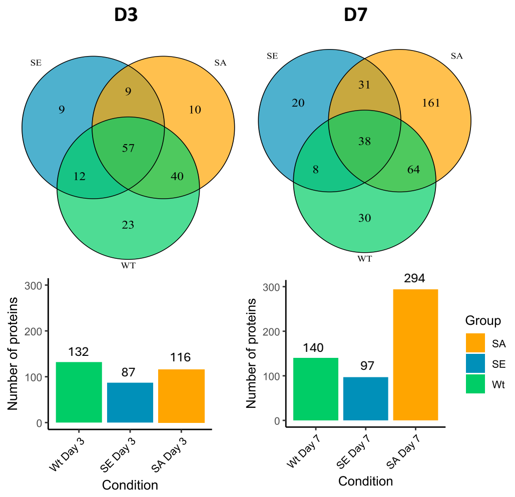


**B.**

| Gene | Accession | PC | Loading | Relative Abundance | | |
|---|---|---|---|---|---|---|
| | | | | APP WT | APP SE | APP SA |
| **PCF11** | O94913 | D3-PC1 | 0.13 | 128.5 | 6.6 | **54.0** |
| **RDX** | P35241 | D3-PC1 | 0.13 | 47.0 | **10.9** | 0.0 |
| **MFAP1** | P55081 | D3-PC1 | 0.13 | 104.2 | 49.6 | **93.2** |
| **YWHAG** | P61981 | D3-PC1 | 0.13 | 44.0 | **23.0** | 0.0 |
| **SF3B6** | Q9Y3B4 | D3-PC1 | 0.13 | 123.9 | **148.6** | 46.9 |
| **NAP1L4** | Q99733 | D3-PC2 | 0.18 | 0.1 | 9.5 | **111.5** |
| **DHX15** | O43143 | D3-PC2 | 0.18 | 133.4 | 21.3 | **111.8** |
| **FXR2** | P51116 | D3-PC2 | 0.17 | 108.3 | **89.2** | 31.7 |
| **JUP** | P14923 | D3-PC2 | 0.17 | 41.5 | 30.8 | **67.3** |
| **PHF5A** | Q7RTV0 | D3-PC2 | 0.16 | 44.3 | **108.8** | 21.3 |
| **SNRPA** | P09012 | D7-PC1 | 0.11 | 159.3 | 68.3 | **128.4** |
| **PRDX1** | Q06830 | D7-PC1 | 0.11 | 1.6 | 0.0 | **193.1** |
| **ENO1** | P06733 | D7-PC1 | 0.11 | 0.4 | 0.0 | **186.6** |
| **DDX17** | Q92841 | D7-PC1 | 0.11 | 78.6 | 29.4 | **118.8** |
| **LMNA** | P02545 | D7-PC1 | 0.11 | 4.8 | 0.0 | **110.1** |
| **FLNA** | P21333 | D7-PC2 | 0.14 | 0.0 | 6.6 | **72.9** |
| **FUS** | P35637 | D7-PC2 | 0.14 | 96.4 | 0.0 | 0.0 |
| **TKT** | P29401 | D7-PC2 | 0.13 | 0.0 | 2.5 | **299.7** |
| **SYMPK** | Q92797 | D7-PC2 | 0.13 | 0.0 | **24.1** | 0.0 |
| **VIM** | P08670 | D7-PC2 | 0.13 | 25.0 | 0.0 | **133.2** |

**Supplementary Figure S1. PCA of the profiles of the APP-GFP samples used, and full number of APP interactors identified by MS if the filter of mandatory presence in 2 replicates is excluded. A.** Principal Component Analysis (PCA) of $log_2$-scaled protein abundances of groups replicates. at day 3 (D3) and day 7 (D7) of differentiation. and all D3+D7 samples together. Profiles based on the data obtained upon imposition of all filters described. APP SA (orange) and WT (green) groups n=2; SE (blue) group n=3. **B.** Table with the top 5 proteins responsible for the groups' separation in each PC1 and PC2 for D3 and D7 of differentiation based on the D3 PCA plot and D7 PCA plot respectively. For PC1 and PC2 at D3, contributes a mix of both (APPSA and APPSE) enriched/exclusive interactors (in bold), while the differences at D7 are greatly driven by interactors exclusive/enriched of the APPSA group. **C.** Venn and bar diagrams of the full numbers of APP interactors identified by MS if the last filter *'only include in a condition the interactors that appear in two replicates of that condition'* is not imposed.

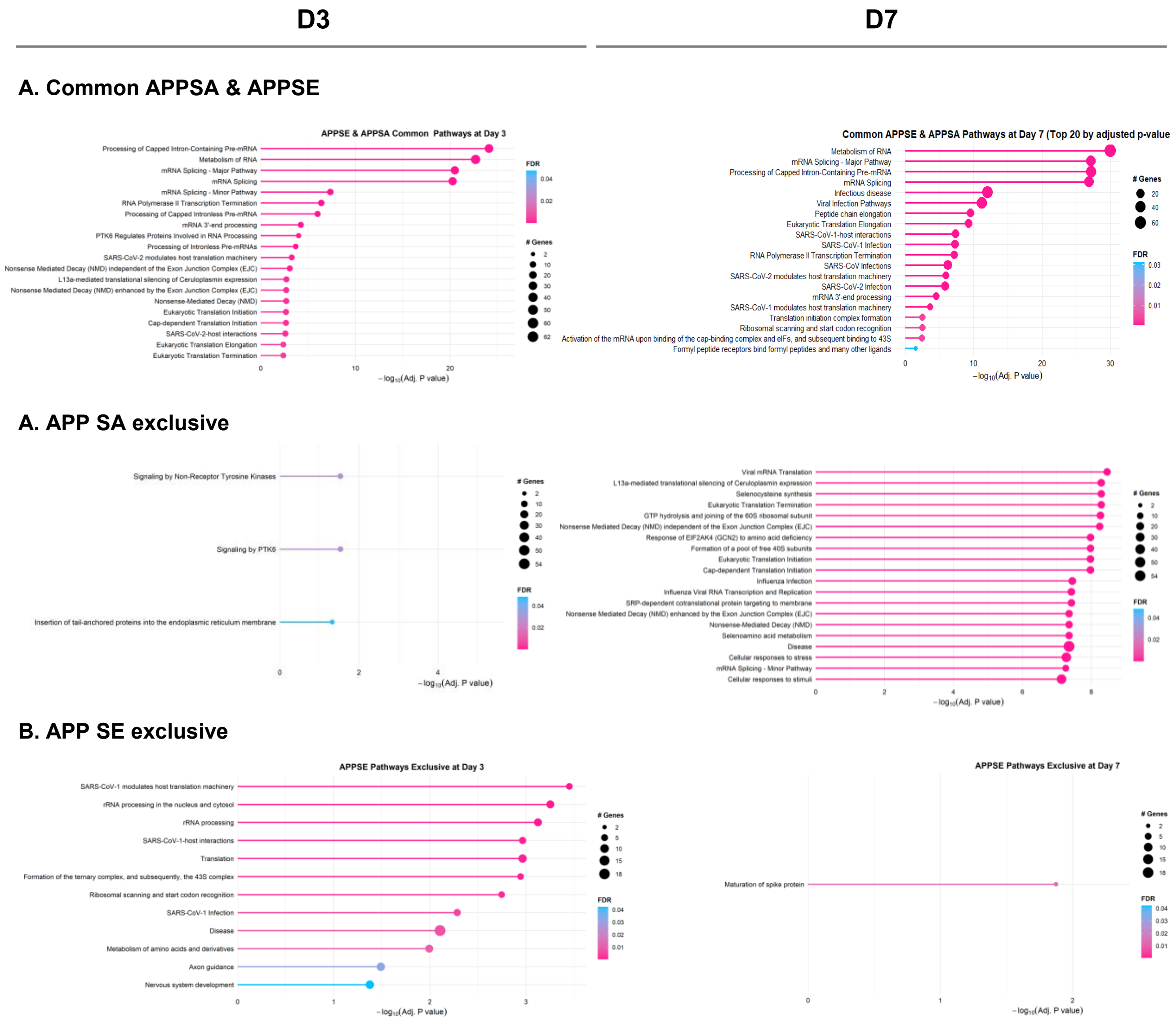


**Supplementary Figure S2. Reactome pathways retrieved using gProfiler with Benjamini-Hochberg p-value correction (p<0.05), ordered by -log10 (adjusted p-value). A.** Pathways common to both APPSA and APPSE interactors lists at D3 or D7 of differentiation. **B.** Pathways exclusively enriched in the APPSA condition, at either D3 or D7. **C.** Pathways exclusively enriched in the APPSE condition, at either D3 or D7. When the number of pathways is superior to 20, only the top 20 are displayed in the plot ordered by adjusted p-value.

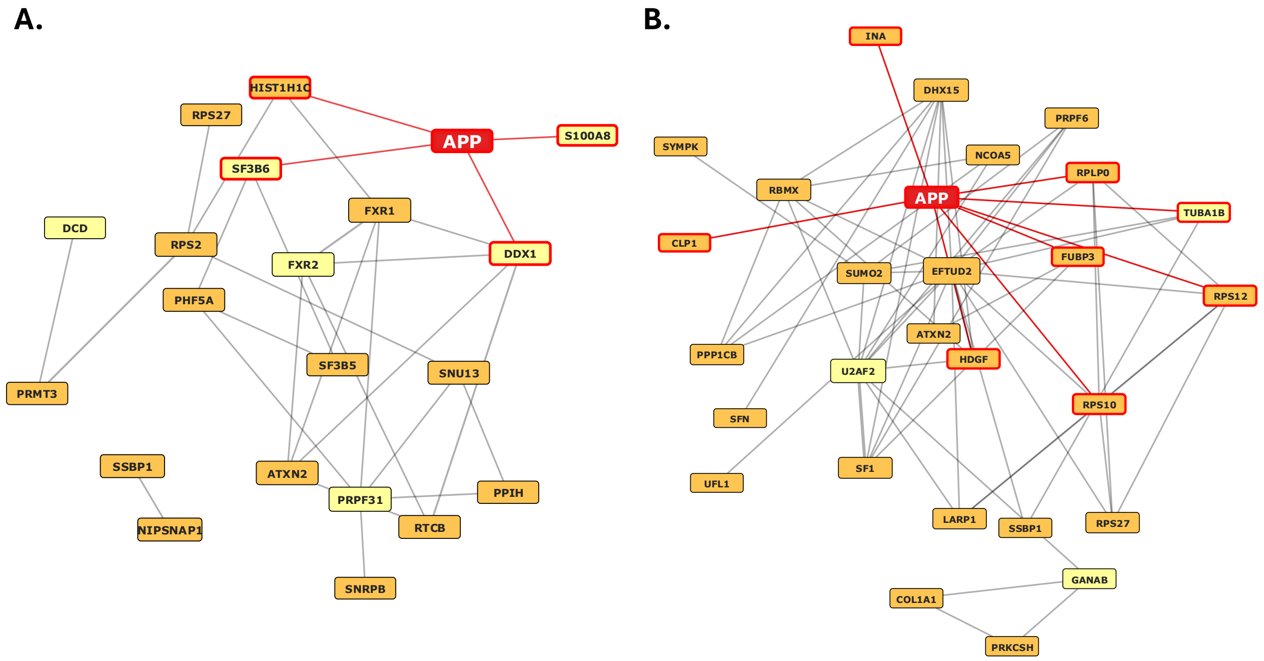


**Supplementary Figure S3. Protein-protein interaction network of the phosphoS655 APP exclusive and enriched interactors at differentiation D3 (A) and D7 (B) if the last filter ('only interactors detected in at least two replicates') was excluded.** Only proteins with node degree > 1 are represented. Node size is based on the node degree and edge width represents the number of experiments supporting the interaction. APP and adjacent edges are represented in red. the nodes connected to those edges have a red outline (interactors already known to interact with APP). Networks built using APID and Cytoscape.

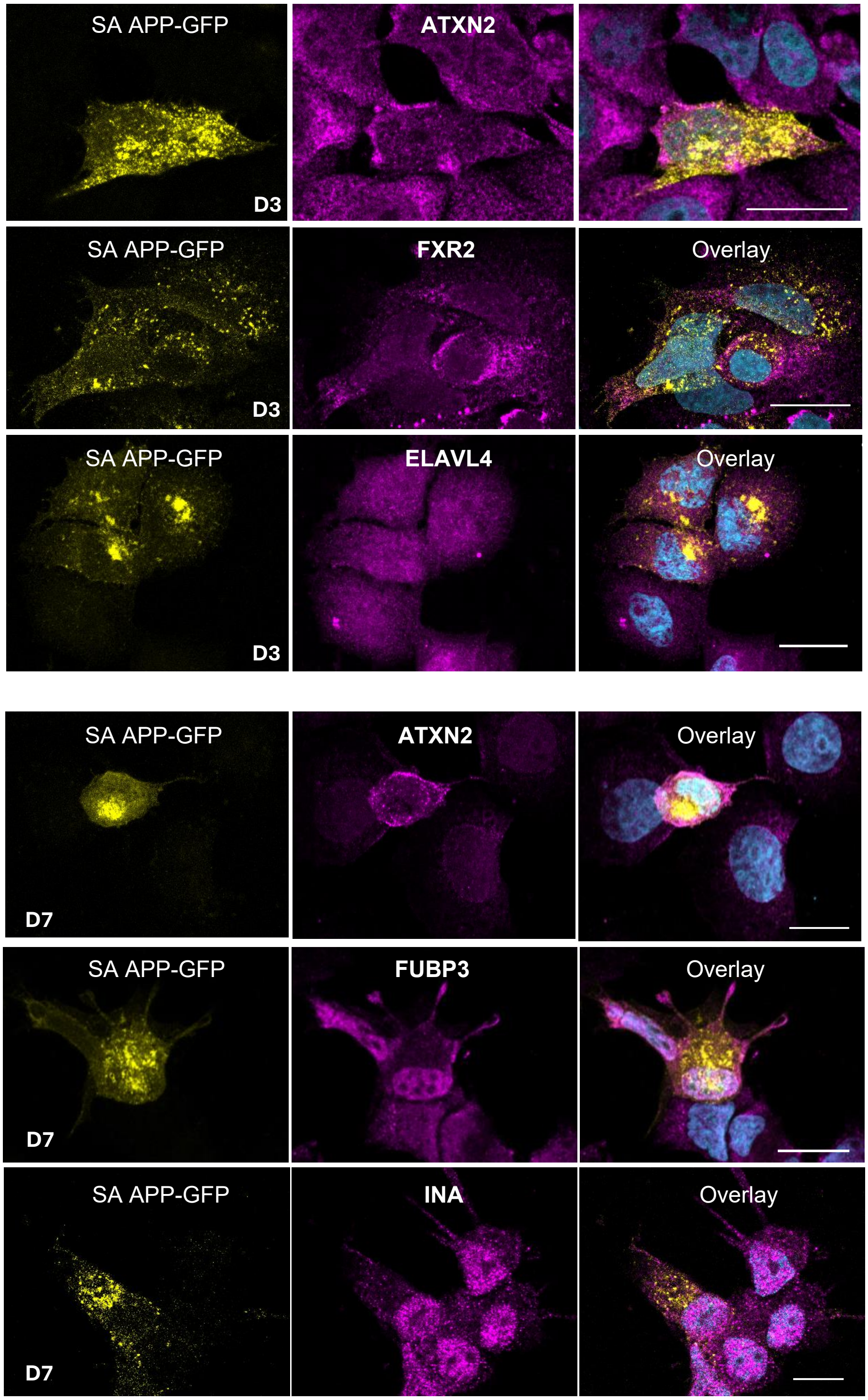


**Supplementary Figure S4. Subcellular distribution of transfected APPSA (SA APP-GFP) and some APPSE-unique or -enriched interactors**, either at day 3 (D3) or day 7 (D7) of SH-SY5Y cells differentiation with retinoic acid. Bar, 20 µm.

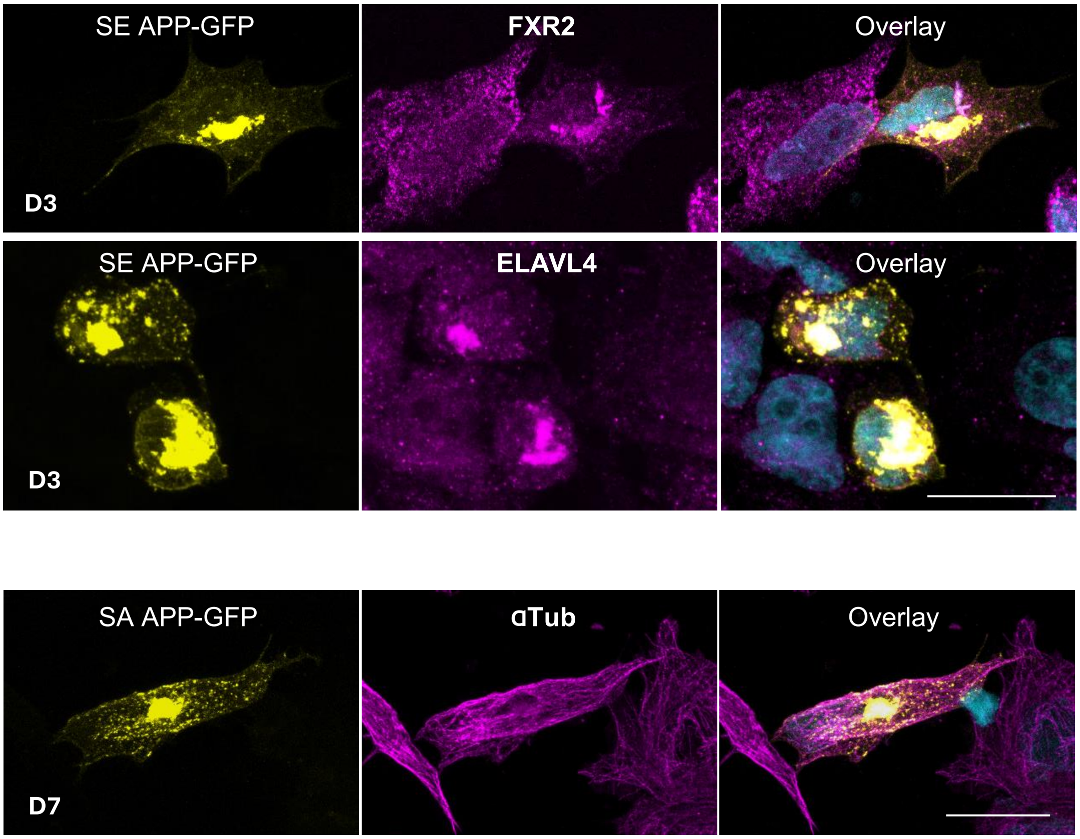


**Supplementary Figure S5. Co-retention of APPSE some enriched interactors in subcellular aggregates when APPSE is highly overexpressed. A.** Immunocytochemistry analysis of the subcellular distribution of APPSE (GFP tagged) and two of its enriched interactors (FXR2 and ELAVL4) in SH-SY5Y cells neuronally differentiated with retinoic acid. Co-retention appears to occur at the Golgi region, strengthening strong interactions between APPSE and these binders. **B.** A third APPSE enriched interactor, alpha-Tubulin, does not appear co-retained with APPSA at the Golfi when this APP form is highly overexpressed at that region. D3/D7, days 3 and 7 of differentiation. Bar, 20 µm.
